# Coumarins disrupt cell-cell communication for control of pathogenesis and virulence in ESKAPEEs and fungal opportunists

**DOI:** 10.1101/2025.04.09.647955

**Authors:** Dylan Boon, Benjamin O’Rourke, Muireann Carmody, David F. Woods, Antje Gloe, Daniel Platero-Rochart, Pedro A. Sánchez-Murcia, Gerard P. McGlacken, F. Jerry Reen

## Abstract

Intricate communication networks and sensing systems underpin the complexity of microbe-host interactions, enabling spatiotemporal control of optimised microbe-host consortia in a diverse range of ecosystems. A central component of these complex interactomes has been the signalling events that enable recognition of host and microbe, whether that niche be clinical or environmental, human or plant. Coumarins have emerged as significant plant derived signalling molecules shaping microbiome dynamics and pathogen behaviours from a broad spectrum of ecosystems. Here we explored the role of natural and synthetic coumarin compounds in signal interference and control of pathogenesis in bacterial and fungal pathogens, uncovering an important ‘hydroxylation-motif’ in the specific inhibition of two *Pseudomonas aeruginosa* interspecies and interkingdom communication molecules. Characterisation of the anti-biofilm activity of coumarins revealed changes in exopolysaccharide production independent of the initial attachment phenotype. Molecular modelling provides an insight into the receptor binding dynamics of three closely related natural coumarins, suggesting an intricate and highly specific host-microbe interaction at the species level. As the very real threat of antimicrobial resistance continues to shadow our horizons, phytochemicals such as coumarins have potential to deliver an ecological solution to dysbiosis in the host-microbe interaction.

**Importance:** Natural ecosystems rely on homeostatic interactions between the kingdoms of life to ensure sustainable and balanced communities can persist. Plant-derived coumarins have recently emerged as playing an important role in shaping microbial communities, presenting a remarkable chemical diversity that can influence the behaviour of bacteria and fungi. At the same time, one of the major challenges to human health continues to be the spread of antimicrobial resistance and the parallel absence of a concerted effort to source and produce new antibiotics at industrial scale. Therefore, new approaches to the control of infection are required, and an ecosystem-level lens may offer one such innovative intervention. Coumarins have the potential to neutralise the very mechanisms used by bacteria and fungi to cause infection leading to morbidity and mortality in hosts ranging from plant to animals. Here we present a mechanistic insight into how effective these molecules can be in targeting keystone pathogens termed the ESKAPEEs and their fungal counterparts.

## Introduction

Important polyphenolic plant metabolites classed as coumarins have recently emerged as key phytochemicals with the potential to shape microbial communities in the rhizosphere (1–5). The manner in which different cultivars of plant host select for specific microbes in the rhizosphere reflects an exquisite level of intricacy in the host-microbe interaction, and in the requirements each distinct cultivar may have for its optimal growth and propagation (6,7). Recently, a mechanistic insight into the coumarin-plant-microbiome relationship has come in the form of MYB72, a host transcription factor and BGLU42, a glucosidase involved in processing the glycoside form of coumarins prior to release from the plant (8). It has also been proposed that mobilisation of iron by plant derived coumarins may underpin, at least in part, the tailoring of the rhizosphere microbiome (1–5,8). However, the extent to which plant-species interactions are governed by coumarin-mediate control of colonisation and polymicrobial interactions remains to be established.

Separately, the growing concern over the inexorable spread of antimicrobial resistance and the parallel drop off in the development of novel antimicrobial compounds has underlined the need for alternative strategies for control of microbial behaviour (9,10). In recent years, there has been a shift towards research into molecules that can intercept the virulence and pathogenic behaviour of microbes, without targeting growth *per se*. The concept of anti-virulence has been tested in a range of bacterial and fungal pathogens, and while some degree of success has been achieved *in vitro* and *in vivo* (11–13), the latter in particular has been with respect to monoclonal antibodies (14–16). The prominent role of coumarins in moderating microbiome dynamics in the rhizosphere suggests that similar structures may be effective in modulating human pathogenic microbes. It remains unclear in this regard whether they act as coercive compounds or rather as cues or signals targeting specific communication systems directly. Elegantly described by Diggle and colleagues, the distinction is an important one (17).

The natural role of coumarins in moderating the behaviour of microbial species led us to explore whether plant-derived coumarins could exert control over key virulence and pathogenesis phenotypes in clinical pathogens, both bacterial and fungal. In particular, we focused on the species of bacteria that pose the greatest risk in the emergence of antimicrobial resistance (AMR), collectively termed ESKAPEE pathogens (18). Previous work has suggested that distinct coumarin molecules can exert control over biofilm formation, cell-cell communication, secretion and growth (19–37), though a universal understanding of the mechanism underpinning these effects remains elusive. Therefore, we explored a broader role for natural coumarins in moderating fungal and bacterial pathogen control, uncovering key structural and mechanistic insights that support a role for coumarins as a plant-derived intervention against fungal opportunists and bacterial ESKAPEEs.

## Results

### Natural coumarins exhibit specific patterns of Quorum Sensing (QS) inhibition in AHL biosensors

Testing the Quorum Sensing Inhibition (QSI) activity of coumarins towards short chain AHL signalling using *S. marcescens* SP19, esculetin (**3**) and esculin hydrate (**4**) exhibited QSI activity at 200, 150, 100, and 50 μg, with the former showing the largest zone of pigment reduction (**Figure 1A** and **Supplementary Figure S1**). Coumarin (**1**), umbelliferone (**2**), and 4-hydroxy-2-coumarin (**5**) also exhibited zones of inhibition at doses of 100 μg and above. In relation to medium-chain AHLs tested on *C. violaceum* DSM30191, coumarin (**1**) and esculetin (**3**) inhibited pigmentation inhibition at 200, 150, and 100 μg. In contrast, esculin hydrate (**4**) showed no inhibition of pigmentation. Inclusion of XTT in the assays provided an assessment of metabolic activity in the zones of clearance, indicating that the reduction in pigmentation in umbelliferone and esculetin was not simply a result of growth inhibition or metabolic suppression (**Figure 1B**). Similarly, coumarin, umbelliferone, and esculetin exhibited activity against the *A. tumefaciens* biosensor, indicating a core set of coumarin structures that may interfere with QS directly (**Figure 1C**).

**Figure 1.**
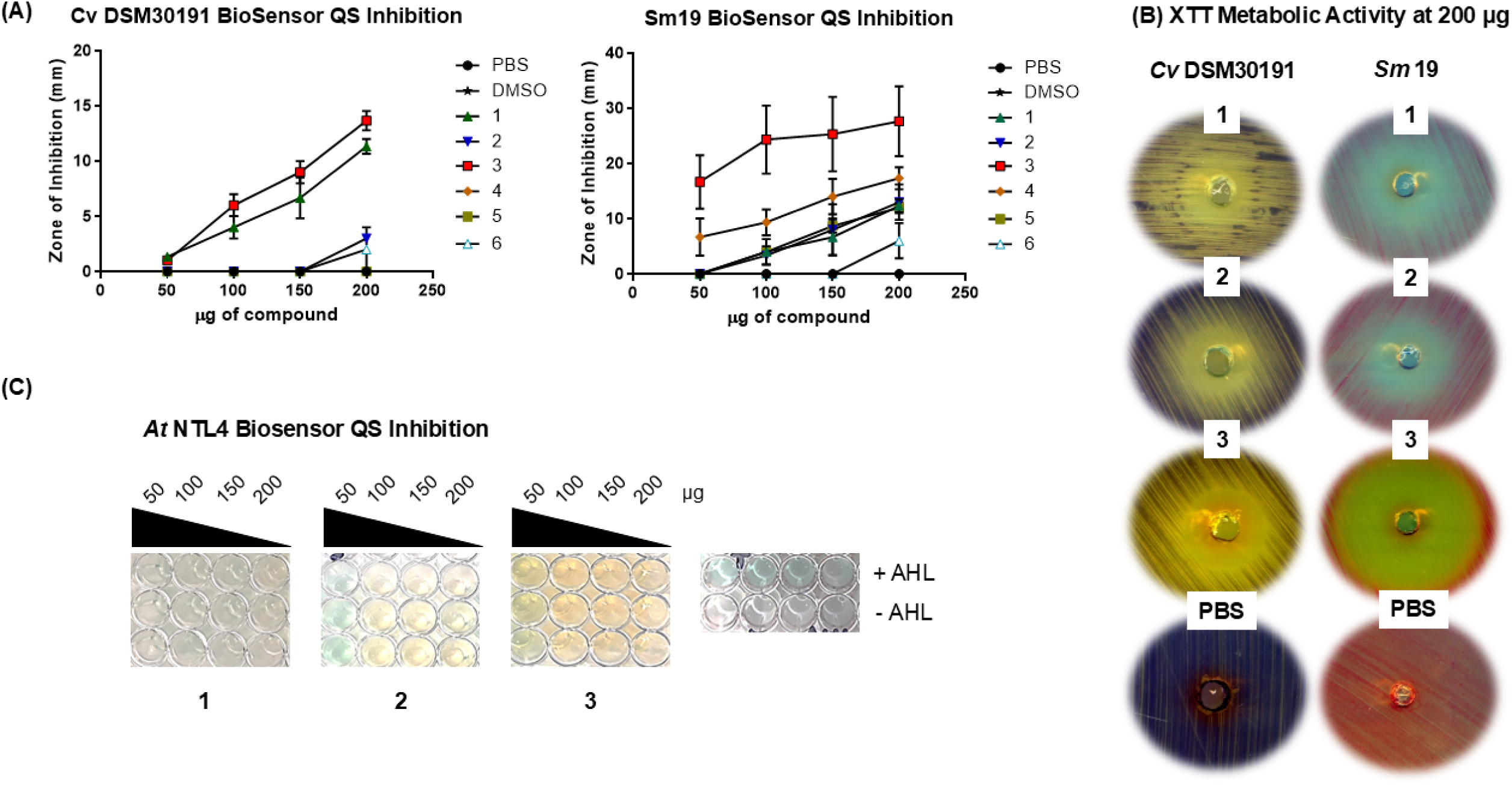
Quorum sensing inhibition by coumarin compounds in AHL-biosensor strains. (**A**) Zone of inhibition from *C. violaceum* and *S. marcescens* biosensors. Data presented is the mean (+/- SEM) of three independent biological replicates. (**B**) Visualisation of XTT metabolic activity in the zone of clearing for the biosensors at 200 μg/ml or coumarin compound. (**C**) Suppression of *A. tumefaciens* NTL4 biosensor by coumarin compounds. Data presented is representative of three independent biological replicates.

### Coumarins suppress biofilm formation in key nosocomial pathogens

Biofilm formation is a major QS-regulated virulence phenotype in pathogens and previous studies have reported anti-biofilm properties of several coumarin related structures. Therefore, the ability of the natural coumarins to interfere with biofilm formation in a range of pathogenic bacteria was investigated. For *P. aeruginosa* PAO1 there was significant antibiofilm properties compared to the DMSO control in coumarin, umbelliferone, esculetin, and 4-methoxy-2-coumarin (**6**) (**Figure 2A**). No significant difference or potential inhibitory effect was seen in the presence of esculin hydrate or 4-hydroxy-2-coumarin.

**Figure 2.**
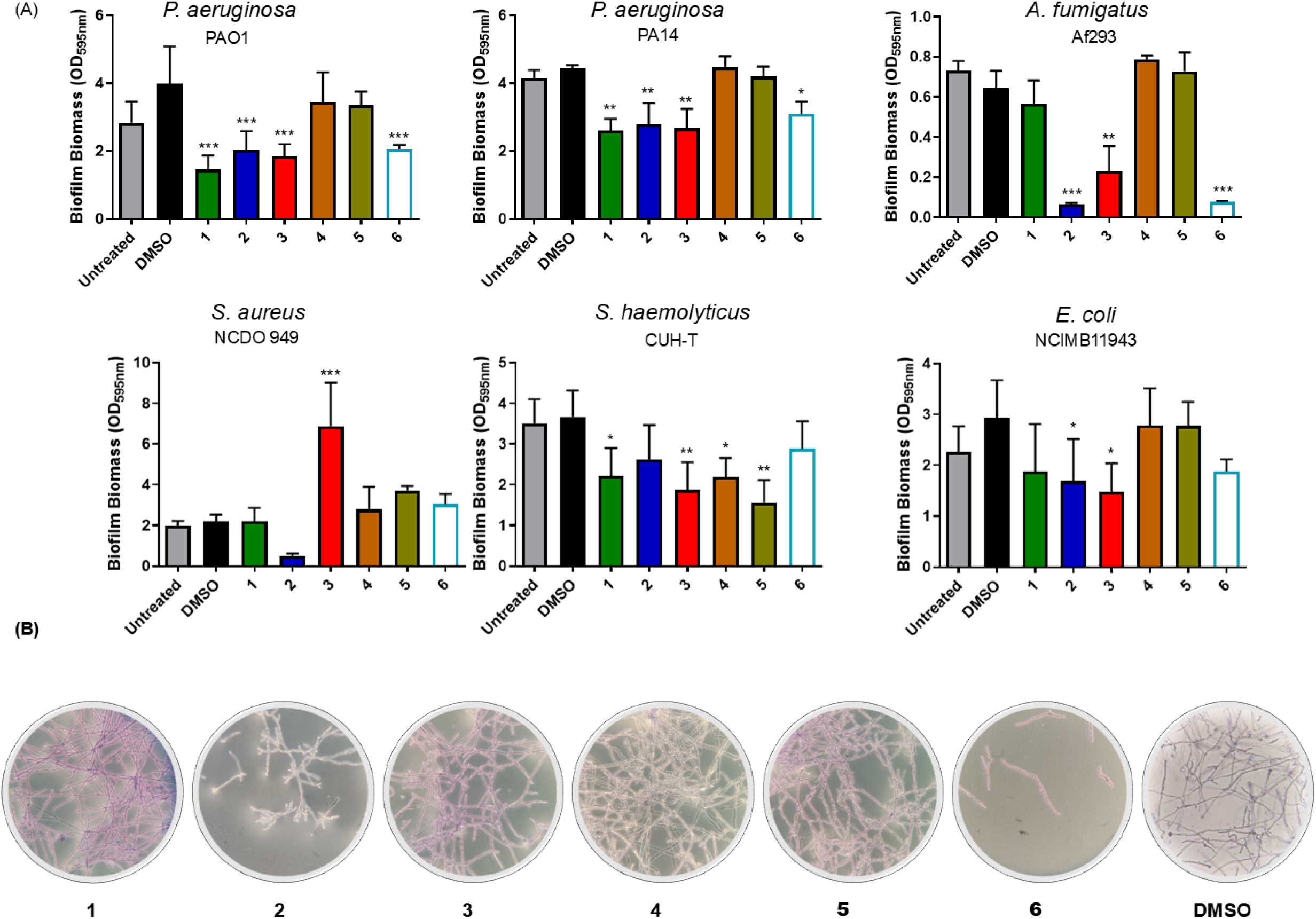
Biofilm formation of key ESKAPEE and fungal pathogens in the presence of coumarin compounds. (**A**) Quantitative analysis of coumarins tested at 2 mM concentration. Data presented is the mean (+/- SEM) of at least three independent biological replicates. Statistical analysis was performed by one-way ANOVA with Dunnett’s post-hoc corrective testing (*p≤0.05, **p≤0.005, ***p≤0.001). (**B**) Microscopic visualisation of hyphal development in A. fumigatus in the presence of coumarin compounds. Data presented is representative of three independent biological replicates.

In *P. aeruginosa* PA14, reduction in biofilm formation was seen in the presence of the same compounds, with the exception of 4-methoxy-2-coumarin. Umbelliferone, esculetin, and 4-methoxy-2-coumarin were also able to suppress biofilm formation in the fungal pathogen *A. fumigatus*, whereas coumarin was ineffective (**Figure 2A,B**). The species-specific nature of coumarin activity was further evidenced by the contrasting activity against biofilm formation in *Staphylococcus* species. Esculetin significantly enhanced biofilm formation in the NCDO949 strain of *S. aureus*, while it inhibited biofilm formation in a clinical *S. haemolyticus* CUH-T strain (**Figure 2A**).

A dose dependent effect was visible in several of the test strains, where at 1 mM coumarin, umbelliferone and esculetin continued to have inhibitory properties on biofilm against *P. aeruginosa* PAO1, while only umbelliferone and esculetin retained activity against the PA14 strain (**Supplementary Figure S2**). No inhibitory effect was evident against any of the tested strains at concentrations of 0.1 or 0.01 µM. MBIC assays determined an IC_50_ of 3 mM for coumarin, 3.2 mM for umbelliferone, and 3.7 for esculetin against *P. aeruginosa* PA14 (**Supplementary Figure S3**).

### Modulation of Exopolysaccharide (EPS) production by specific natural coumarins

Biofilm formation is a multi-stage process and interference with it can occur at distinct phases. The *P. aeruginosa* PA14 strain was selected for analysis based on its strong biofilm-forming properties and its well characterised EPS morphology. Initial attachment was unaffected indicating that inhibition of biofilm biomass occurs downstream of microcolony formation (**Figure 3A**). Assessing biofilm formation over time, it became apparent that the suppression phenotype occurs between 2-4 hrs post inoculation, with a significant increase in biofilm biomass occurring in the carrier control, where biofilm biomass remained unchanged in the coumarin treated samples (**Figure 3B**). We next studied production of exopolysaccharide to determine if EPS was the target of coumarins, thus preventing the mature biofilm from forming (**Figure 3C** and **Supplementary Figure S4**). EPS production appeared comparable to the DMSO control in the presence of esculetin. However, pigmentation and morphology appeared different in the samples treated with coumarin and umbelliferone, with a more intense red colour than observed in the DMSO control, comparable to the *tbpA-D* mutant.

**Figure 3.**
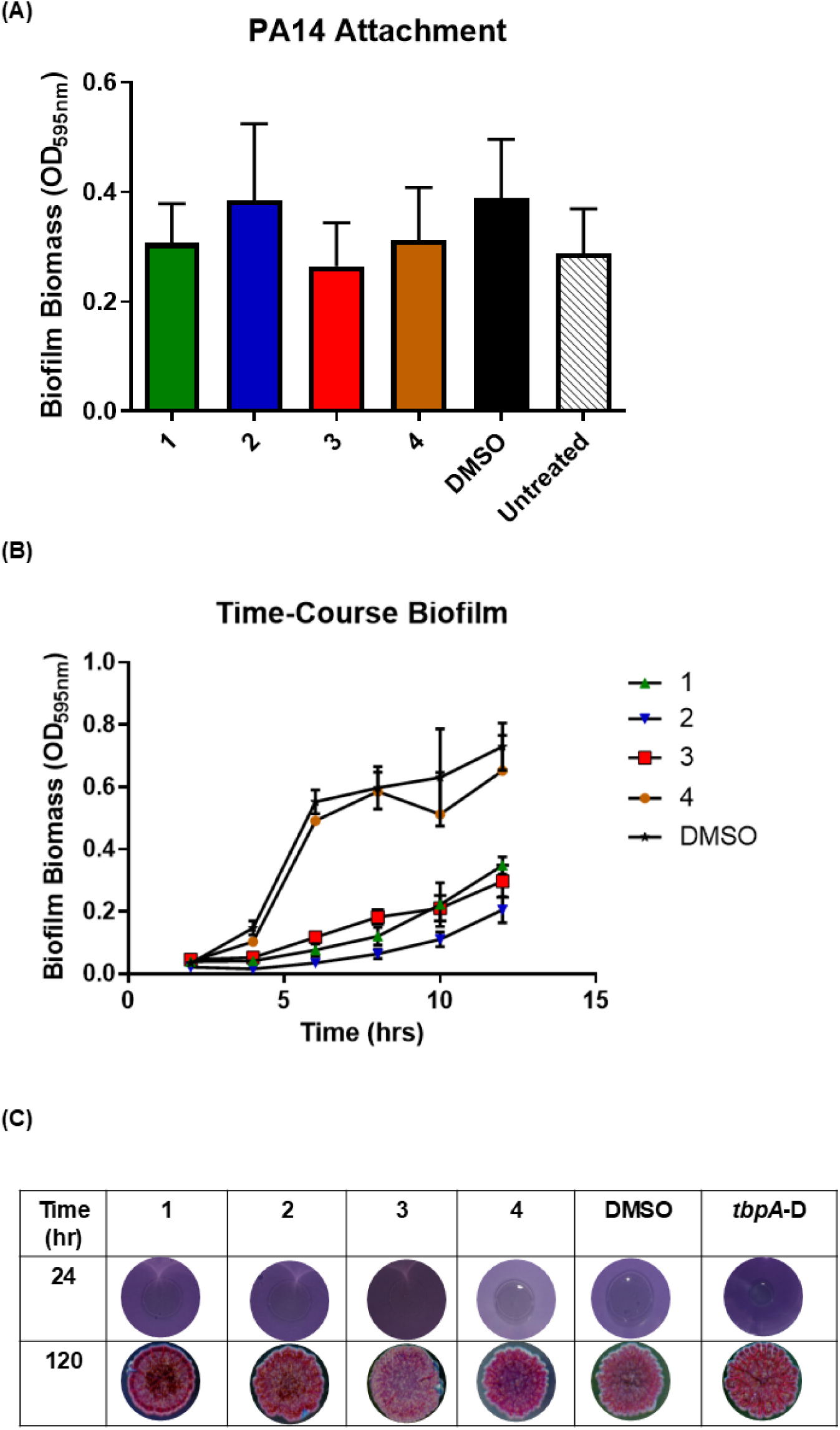
Biofilm (attachment and time-scale) analysis of *P. aeruginosa* PA14 in the presence of selected coumarin compounds. (**A**) Attachment analysis performed over 2 hrs on microtitre plates. (**B**) Time-course biofilm kinetics measuring biomass over a 10 hr period from the point of inoculation. (**C**) EPS production measured over 120 hrs in the presence of selected coumarin compounds. All data presented is the mean (+/- SEM) of at least three independent biological replicates. Statistical analysis was performed by one-way ANOVA with Dunnett’s post-hoc corrective testing.

Growth was also investigated, as a growth limiting activity could also explain the biofilm results. *P. aeruginosa* PAO1 and *S. haemolyticus* CUH-T had no significant differences in growth when treated with the coumarin compounds (**Supplementary Figure S5**). *P. aeruginosa* PA14 entered stationary phase at a lower biomass in the presence of coumarin, umbelliferone and esculetin, while in *S. aureus* NCDO949 coumarin led to a slower growth rate but same final biomass comparative to the DMSO control. For *E. coli* NCIMB11943, there was significant growth differences for the bacteria inoculated in the presence of coumarin, umbelliferone and esculetin. While these growth effects may explain the reduction in biofilm formation for *E. coli,* they indicate that the suppression of biofilm formation in ESKAPE pathogens by coumarins is generally a growth independent phenotype.

### *P. aeruginosa* QS systems are suppressed by specific coumarin compounds

Previously, coumarin has been shown to suppress the LasIR, RhlIR, and PQS QS signalling systems (21,23,25,33). However, the mechanism of action and the role of other coumarin compounds in this regard remains uncharacterised. Promoter activity was initially studied at mid-log and early stationary phase timepoints (**Figure 4A**). Interestingly, esculetin led to increased activity at the *lasI* promoter, counterintuitive to what might be expected from the biosensor assays. Both coumarin and umbelliferone led to a significant suppression of *pqsA* promoter activity in the wild-type strain, while esculetin and the other coumarins did not influence activity relative to the DMSO control (**Figure 4B**). Addition of coumarins did not affect promoter activity of *rhlI*, again surprising given that it suppressed QS signalling in the *S. marcescens* S19 biosensor (**Figure 4C**). The activity of coumarin, umbelliferone, and esculetin was subsequently assayed over time-course in shaking flasks to conduct a more detailed kinetic analysis of promoter activity from the QS systems. This confirmed the enhanced *lasI* promoter activity in the presence of esculetin (**Figure 4D**), and the suppression of *pqsA* promoter activity by coumarin and umbelliferone (**Figure 4E**). No significant change in *rhlI* promoter activity was observed at any of the time points tested (**Figure 4F**).

**Figure 4.**
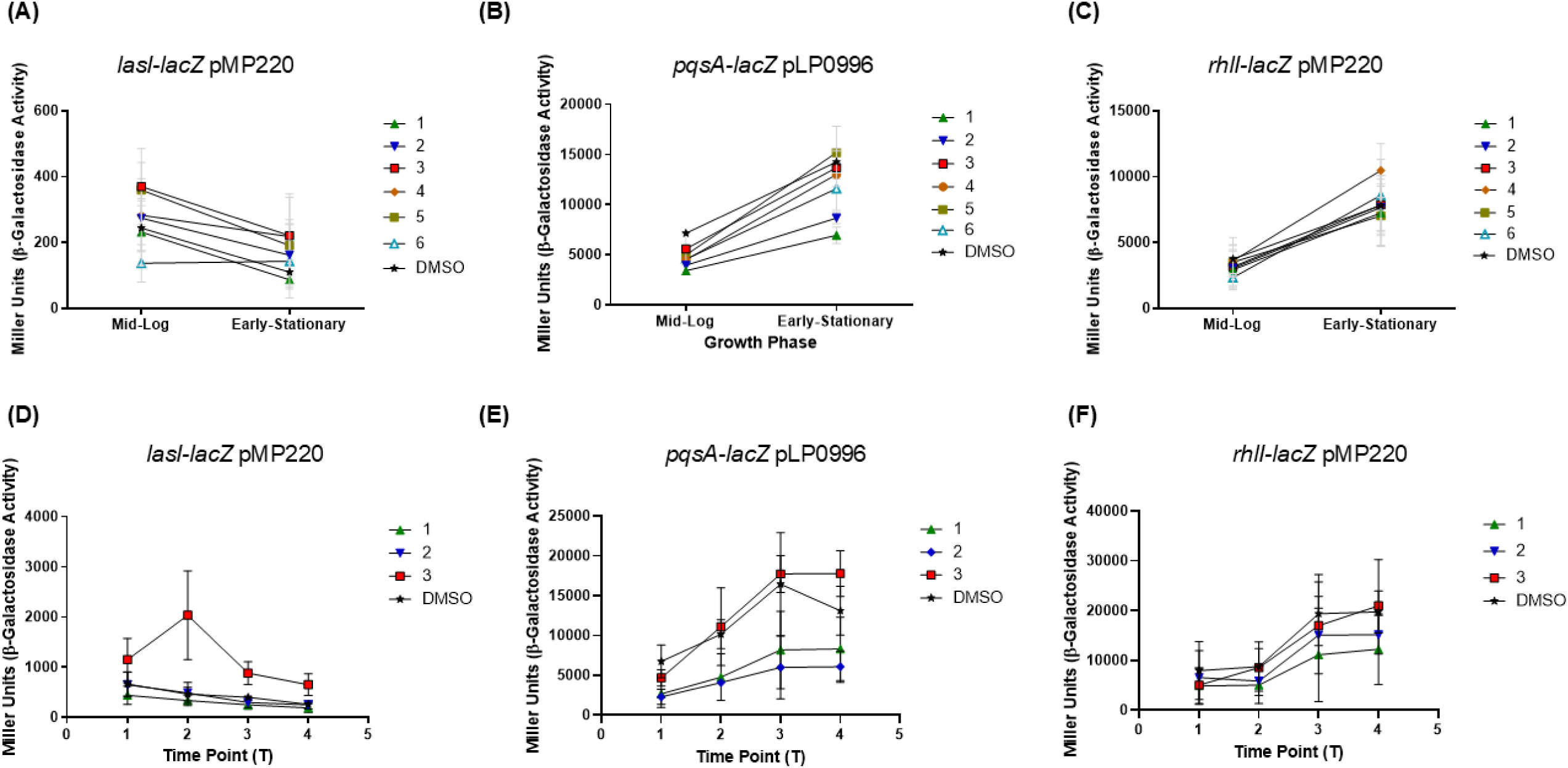
Quorum sensing promoter fusion analysis in *P. aeruginosa*. The three QS systems encoded in *P. aeruginosa* were studied in the presence of coumarin compounds. (**A-C**) Assays performed in glass vials and (**D-F**) time-course assays performed in conical flasks. All data presented is the mean (+/- SEM) of at least three independent biological replicates.

The suppression of *pqsA* promoter activity by coumarin and umbelliferone was of particular interest in light of the role played by HHQ and PQS in interkingdom communication and the host-pathogen interaction (38). These two quorum sensing signals have previously been shown to operate through the PqsR (also known as MvfR) LysR-Type transcriptional regulator and have exhibited interspecies and interkingdom properties in several studies (38–41). To further explore the impact of these natural coumarin compounds on *pqsA* promoter activity, the impact of coumarins on activation of the system by exogenous PQS was explored using a *P. aeruginosa pqsA* mutant carrying the pLP0996 reporter. Addition of exogenous PQS led to a significant increase in *pqsA* promoter activity, thus serving as a positive control (**Figure 5A**). When exogenous PQS was added in the presence of coumarin compounds, coumarin and umbelliferone both blocked the activation of the *pqsA* promoter, suggesting that suppression of *pqsA* promoter activity may be the result of a direct influence on the PqsR protein (**Figure 5A**). The suppression of *pqsA* promoter activity is the result of competitive inhibition, with dose response curves confirming the shift in suppression at lower doses of coumarin and umbelliferone (**Figure 5B**). Promoter fusion analysis of a *pqsR* translational fusion indicates that *pqsR* promoter activity is itself reduced in the presence of coumarin and umbelliferone, raising the possibility that auto-induction may be impaired, or that a secondary target for coumarins could suppress activation at this locus (**Figure 5C**). The suppression of *pqsA* promoter activity would be expected to result in a reduction in HHQ and PQS signal production in the presence of coumarin and umbelliferone. This appeared to be consistent with TLC analysis. Extraction of samples grown in the presence of esculetin appeared to result in higher signal production than the DMSO control. Production of the virulence factor pyocyanin, which is regulated by the PQS system in *P. aeruginosa*, was decreased in the samples grown with coumarin and umbelliferone when compared to the DMSO control (**Figure 5E**), consistent with a loss of PQS production.

**Figure 5.**
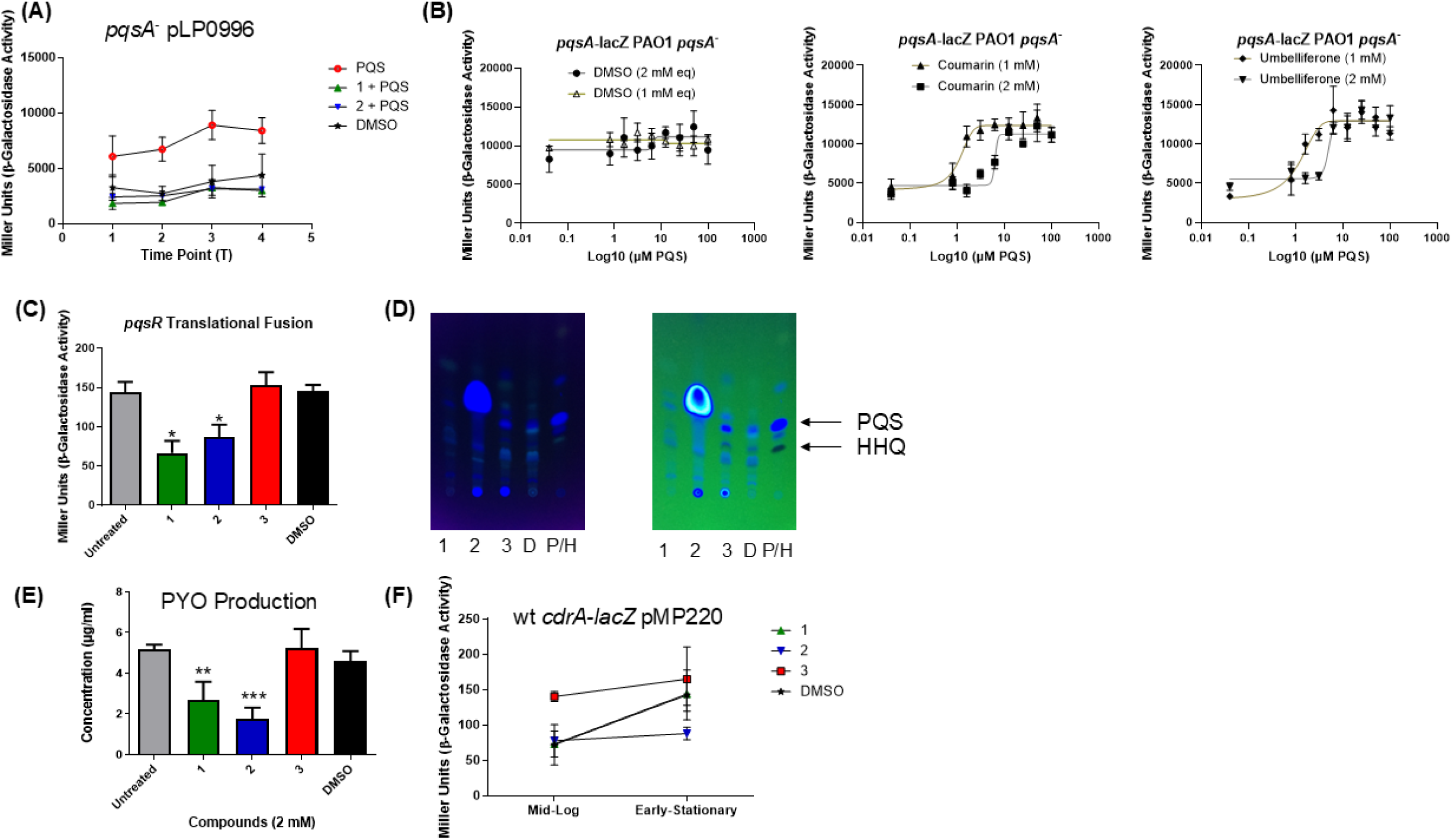
Direct QS promoter activity suppression in a *pqsA* mutant. (**A**) *pqsA* promoter activity in a *pqsA* mutant grown in the presence of coumarin compounds in the presence of exogenous PQS. (**B**) Competitive inhibition analysis of the coumarins vs PQS PqsR interaction using a *pqsA* mutant *pqsA* promoter fusion with increasing concentrations of PQS. (**C**) *pqsR* promoter activity in the wild-type *P. aeruginosa* strain in the presence of coumarin compounds. (**D**) TLC analysis of *P. aeruginosa* extracts from growth in the presence of coumarin compounds. Images are representative of at least three independent biological replicates. (**E**) PYO production and (**F**) second messenger signalling in *P. aeruginosa* in the presence of coumarin compounds. All data presented is the mean (+/- SEM) of at least three independent biological replicates. Statistical analysis was performed by one-way ANOVA with Dunnett’s post-hoc corrective testing ((*p≤0.05, **p≤0.005, ***p≤0.001).

Analysis of the impact of coumarin compounds on cyclic-di-GMP second messenger signalling indicated that umbelliferone alone suppressed promoter activity of the *cdrA* promoter biosensor (**Figure 5E**). While coumarin itself has previously been shown to reduce cyclic-di-GMP levels in *P. aeruginosa* at the concentration tested (24), the lack of activity against the *cdrA* promoter observed here suggests its influence may occur independent of the CdrA network.

### Molecular modelling reveals structural insights into coumarin species specificity

To investigate further how coumarin (**1**) and umbelliferone (**2**) could interfere with cell-cell communication in *P*. *aeruginosa*, we explored the possibility that these molecules could block either the interaction between the autoinducer N-3-oxo-dodecanoyl-L-homoserine lactone (3-oxo-C12-HSL) and the transcriptional activator protein (LasR) and/or the interactions between PQS and the LysR-type transcriptional regulator PqsR. Esculetin (**3**), which contains hydroxyl substitutions at both the 6-, and 7-, positions was included as a reference point, in light of its inactivity against PQS signalling.

First, we docked the three molecules at the binding sites found for PQS (PDB id. 6YIZ) (42) and 3-oxo-C12-HSL (PDB id. 2UV0) (43) in PqsR and LasR, respectively (**Figure 6A**). No significant differences were found between the pose solutions of the three molecules to both proteins. Thus, we considered the docking solutions as initial conditions of a short Molecular Dynamics (MD) simulation in order to identify the fingerprint of the main interactions of coumarin and its two hydroxy-derivatives with LasR and PqsR in the docked binding sites. In PsqR, esculetin barely changes its orientation obtained by the docking studies. In contrast, coumarin and umbelliferone show RMSD values above 4 Å. Much smaller RMSD variations were observed when bound to LasR (**Figure 6B**).

**Figure 6.**
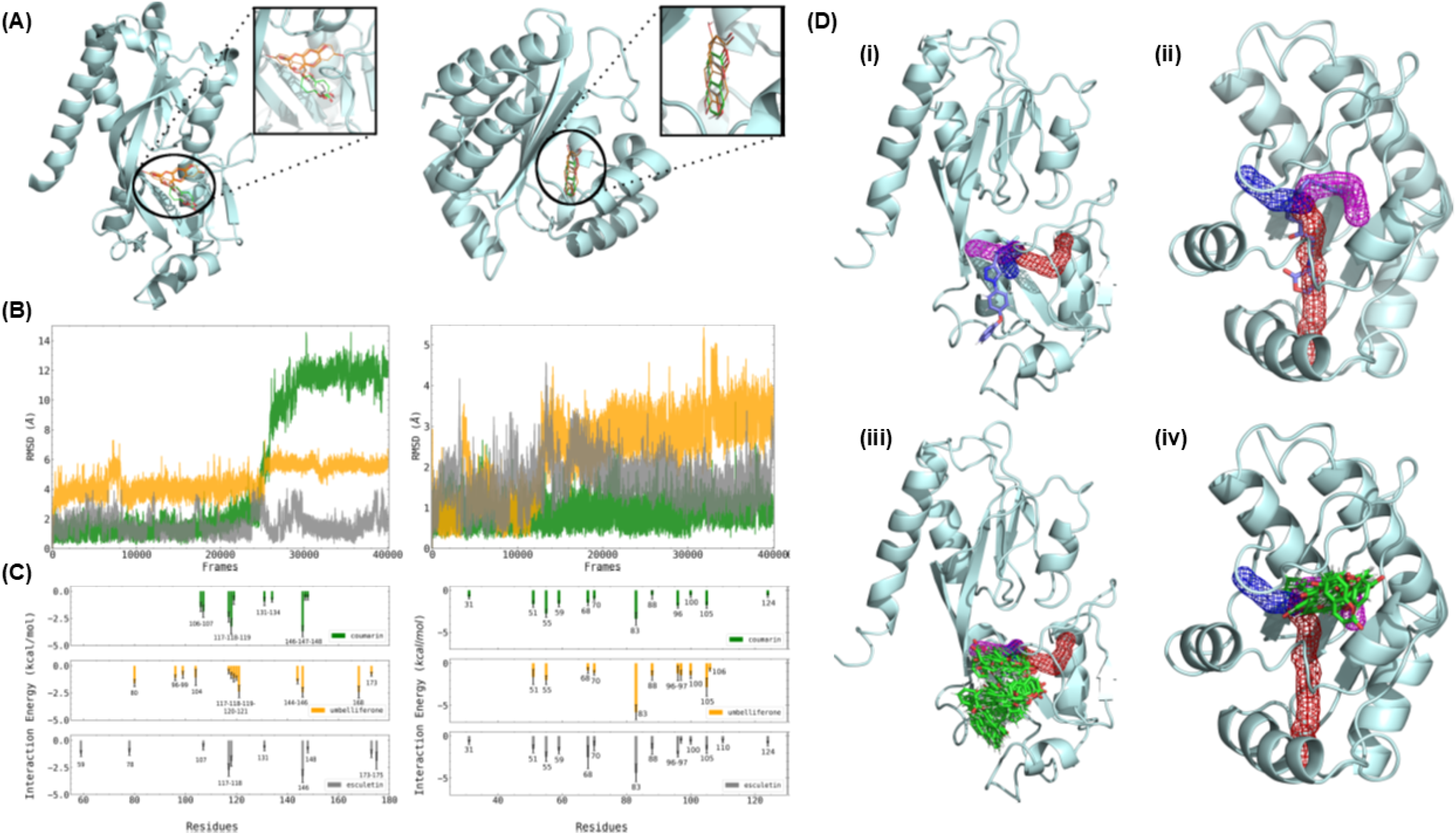
Molecular modelling of the PqsR and LasR interaction with coumarin, umbelliferone, and esculetin. (A) Representative docking poses obtained for coumarin (C-atoms in green), umbelliferone (orange) and esculetin (grey) in PqsR (left) and in LasR (right). (B) Root-mean square deviation (RMSD, Å) for coumarin (green line), umbelliferone (orange) and esculetin (grey) bound to PqsR (left) and LasR (right). (C) Interaction ligand-protein energy (kcal mol −1) decompose per residue for the receptors PqsR (left) and LasR (right). Error bars count for the standard deviations (30 snapshots in a 30ns-window from the MD simulation). (D) Superposition of the apo structure of PqsR (i) and LasR (ii) and the crystal structure with the co-crystallised ligand (represented as sticks in purple). In the mesh representation, tunnels 1 (blue), 2 (magenta), and 3 (red) found by the Caver are depicted. Tunnels explored by 2 during the MD simulations are presented for PqsR (iii) and LasR (iv).

The interaction fingerprints for the three compounds bound to both receptors PqsR and LasR were computed via the algorithm MM-ISMSA3 (**Figure 6C**). In general, the interactions observed in PqsR are mainly weak dispersion interactions. As an example, the C_sp_^3^H-π interaction between the methyl groups of Ile146 in PqsR with the electron-deficient pyrone ring of the three coumarin ligands is the one with the larger contribution. The interactions of these molecules with other aliphatic residues like Leu117 is also notable. Furthermore, the carbonyl oxygen in the pyrone ring points to the region where residues Leu106 and Ser107 are located and forms temporal hydrogen bonds with their hydroxyl and amino groups, respectively. Interestingly, the hydroxyl group on position 7-(umbelliferone) and on both positions 6- and 7-(esculetin), do present new hydrogen bonds for the protein-ligand interaction. In the case of umbelliferone, a hydrogen bond is found between the hydroxyl group and the aromatic ring of Tyr168, while the carbonyl oxygen (in the pyrone ring) establishes a hydrogen bond interaction with the side chain of Gln104. In the case of esculetin, the hydroxylation on C-6 introduces a hydrogen bond with residue Thr175, with the 7-OH group projecting into the solvent. Regarding LasR, aromatic residues like Trp83 and Trp55 are prominent in the binding site. Indeed, Trp83 presents the largest contribution for the energy binding of the three molecules. Finally, the global binding energy (**Supplementary Table S1**) corroborates that there are also no appreciable differences amongst the three molecules in terms of their energetic potential.

As a second strategy, we ran longer MD simulations using the apo form of the proteins embedded in an aqueous solution including 10 molecules of each of the coumarin derivatives per protein chain. This enabled us to check if the molecules in solution could enter the former binding sites explored in our docking studies. In the case of LasR we decided to use a dimer since this is the biologically relevant form as described by Bottomley and colleagues As can be seen from the RMSD evolution along the MD simulation in **Supplementary Figure S6**, at least one molecule of coumarin, umbelliferone and esculetin is able to approach and enter to the binding site in PqsR (RMSD values highlighted with a dotted-line box). On the other hand, none of the molecules appear to reach the active site in the case of LasR (**Supplementary Figure S6**). In order to understand the differential behaviour of the two proteins in our MD simulations regarding the access of the ligands to the crystallographic binding site, we analysed relaxed structures of the apo form of both proteins using the software Caver (44). In PqsR, the co-crystallised triazolopyridine inverse agonist A overlaps mainly with the magenta and blue tunnels; in LasR, the overlap of 3-oxo-C12-HSL is mainly with the red and the magenta tunnels (**Figure 6D(i** and **iii)**). Comparison of the solutions of the MD simulation for coumarin revealed that whereas in PqsR coumarin is able to explore the two tunnels where the co-crystallised ligand is present, in LasR the corresponding one (red tunnel) is not explored (**Figure 6D(ii** and **iv)**). Calculating the size of the bottleneck radius (Å) and the average throughput for the six tunnels (**Supplementary Table S2**), the blue tunnels were found to be the most probable for molecule transit.

### Structure function profiling indicates a core coumarin framework for species control

Evidence of structural differences underpinning the biological activity of the coumarin compounds was apparent in the phenotypic studies with hydroxylation (comparing compounds **1-3**) and the hydroxyl vs methoxy group at position 4 (comparing compounds **5-6**) resulting in markedly different phenotypes in ESKAPE pathogens. To explore this further, a total of four coumarin analogues (**7**-**10**) were tested, including warfarin (**7**) and a fluorescent probe (**8**). The other two compounds were analogues of the pyrone component of the coumarin structure (compounds **9-10**). With the exception of warfarin leading to a slower growth rate and lower final biomass in *S. aureus* NCDO949, growth was uninhibited by the coumarin analogues (**Supplementary Figure S7**).

None of the synthetic analogues exhibited suppression of pigmentation in the *C. violaceum* DSM30191 or *S. marcescens* Sm19 biosensors (**Supplementary Figure S8** and **S9**). The *A. tumefaciens* biosensor appeared more susceptible to modified coumarin challenge, with compound **10** (4-hydroxy-6-methyl-2-pyrone) exhibiting activity. While none of the modified coumarin compounds exhibited anti-biofilm activity against *P. aeruginosa*, compound **8** was notable for its biofilm inhibitory activity towards *A. fumigatus*, *Staphylococcus*, and *E. coli*, while **9** was also suppressive of biofilm formation in these species with the exception of *E. coli* (**Figure 7** and **Supplementary Files S10-S12**). It was noteworthy that while **7** and **9** significantly enhanced biofilm formation in *S. aureus* NCDO949, they had a strong suppressive effect against *S. haemolyticus* CUH-T. From the perspective of the three QS systems in *P. aeruginosa*, compound **7** led to increased *lasI* promoter activity, while **9** resulted in reduced *lasI* and *pqsA* activity (**Figure 7** and **Supplementary Figure S13**). Compounds **8** and **10** did not influence promoter fusion activity (**Figure 7** and **Supplementary Figures S13**). It was also notable that compound **9** led to a reduction in PYO production in *P. aeruginosa*. Finally, compounds **7** and **9** increased *cdrA* promoter activity in mid-log phase (**Figure 7**).

**Figure 7.**
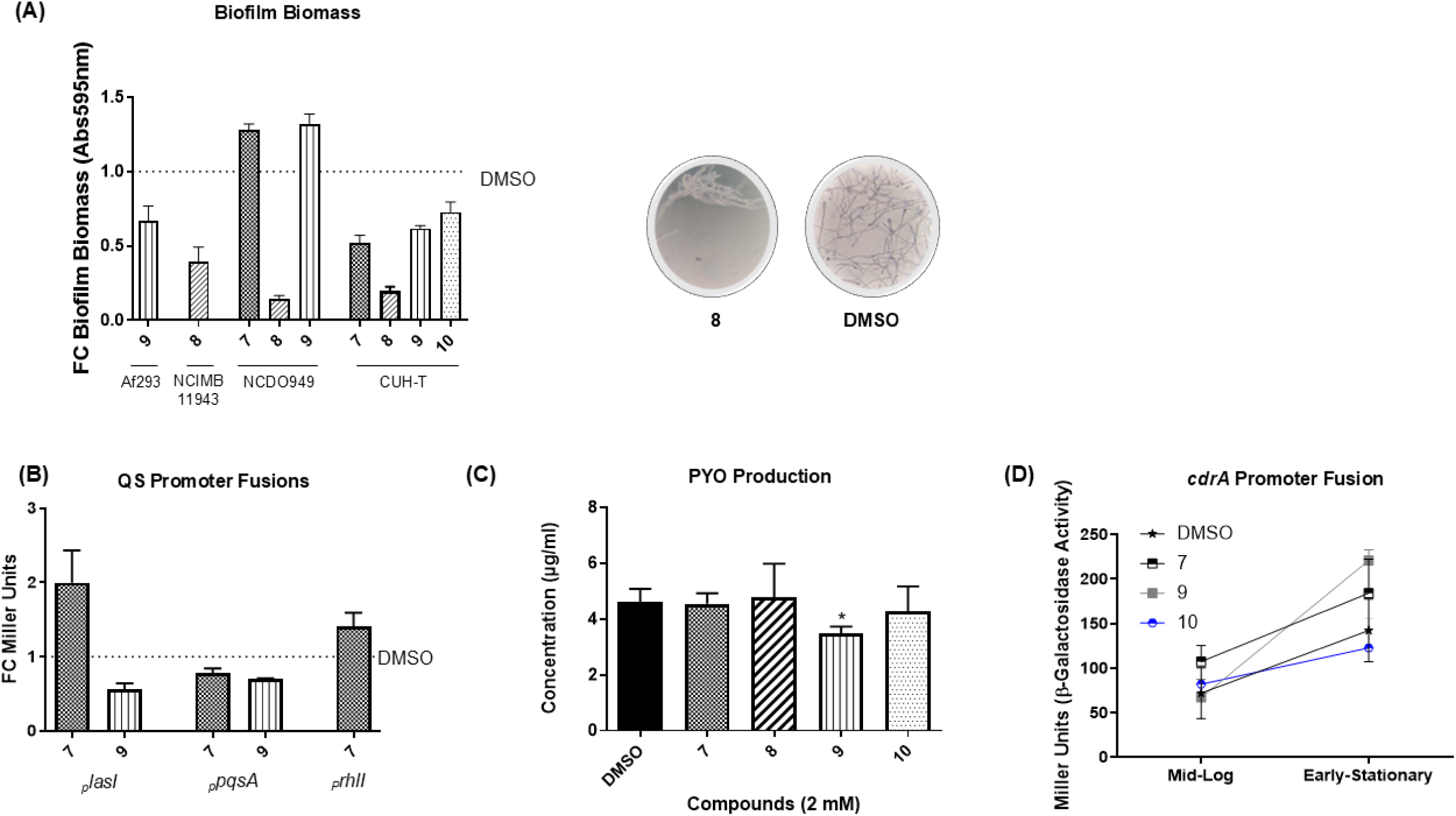
Profile of activity of synthetic coumarin compounds against key pathogens. (**A**) Fold change in biofilm formation and visualisation of *A. fumigatus* Af293 hyphal structures; (**B**) Fold change in promoter activity for QS systems (**C**) PYO production in *P. aeruginosa* PAO1, and (**D**) *cdrA* promoter activity are presented. All data presented is the mean (+/- SEM) of at least three independent biological replicates.

## Discussion

The increasing prevalence of antibiotic resistance, especially in nosocomial infections, is becoming a serious global threat to public safety (45). Apart from the well documented projected increased mortality arising from AMR, illnesses due to antibiotic resistant bacteria may also take longer to resolve and will increase healthcare expenses (46). The dearth of antibiotics in development has led to studies on alternative treatment strategies, such as antimicrobial molecules and compounds that can target specific virulence factors and communication systems, i.e. quorum sensing (13,23,41,47–51). Though the view was long held that targeting of QS poses a lower risk of resistance developing, there is growing evidence that resistance has been encountered in laboratory and clinical strains (52,53). While it is likely that a less potent ‘Darwinian’ selection may manifest in the presence of these interventions, the design and future development of small molecular approaches to pathogen control must heed these concerns.

Coumarins are natural plant secondary metabolites that may have a key role in targeting microbial communication systems (**Figure 8**). Coumarin compounds have been explored as potential combinatorial adjuncts for antibiotic treatments, with some evidence suggesting synergistic effects observed against a range of multidrug resistant pathogens (54–56). The mechanism of action underlying antibacterial activity or antibiotic potentiation has not yet been uncovered, though possible roles for quorum sensing, dihydrofolate reductase, and efflux have all been reported (21,24,25,33,55,57). Apart from their emerging role in shaping the rhizosphere, or perhaps as a consequence of this, coumarins present a significant opportunity as a framework for the design of therapeutic interventions against opportunistic pathogens. The coumarin family of compounds have shown attractive pharmacological characteristics such as anti QS, antibiofilm, antifungal and antioxidant properties (23,58–61). Coumarins have also demonstrated repression of the classic AHL, and the more species-restricted alkyl-hydroxyquinolone (AHQ) quorum sensing systems (21,23–25,33).

**Figure 8.**
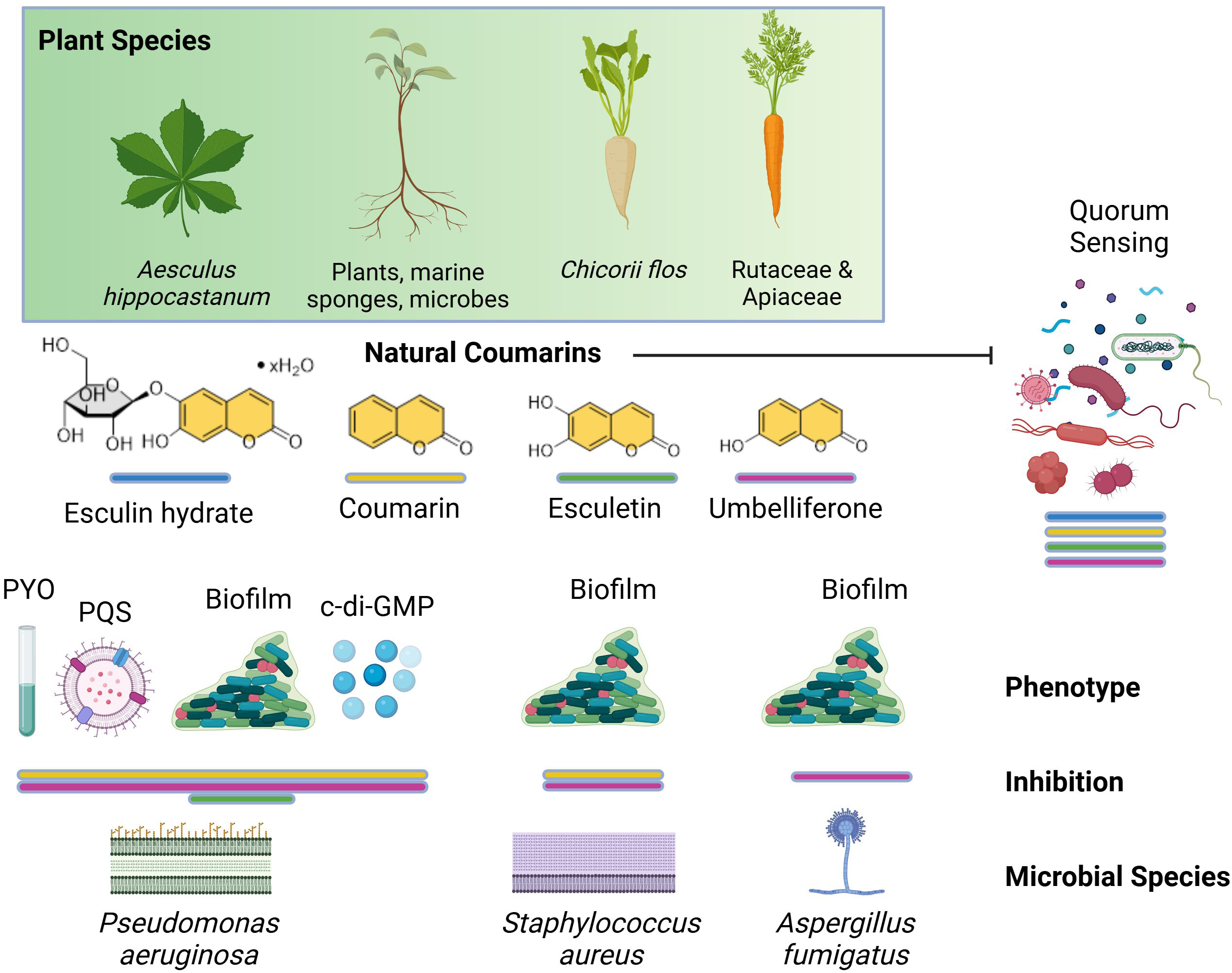
Overview of coumarin related anti-infective traits and the species specificity with which plants can influence microbial behaviour. The selective nature through which coumarin compounds produced by different species of plants select for distinct phenotypic responses in key pathogenic organisms adds to the growing evidence of cultivar-mediated microbiome shaping as a hallmark of the plant-microbiome interactome in the rhizosphere and marine ecosystem. Created with BioRender.com.

The importance of the hydroxyl group with respect to QS inhibition uncovered in this study is consistent with previous findings by D’Almeida and colleagues where coumarin molecules with hydroxyl groups on the aromatic ring were found to display enhanced inhibition of biofilm formation by *P. aeruginosa* when compared with coumarins with substituents in positions 3 and 4 or without the double 3,4 bond (20). The degree of hydroxylation and/or position of the hydroxy groups has a significant impact on the anti-infective activity. QS and biofilm formation were inhibited by coumarin (no hydroxy group), umbelliferone (7-hydroxy) and esculetin (6,7-dihydroxy) in several pathogenic species. Esculetin exhibited the broadest spectrum of activity, targeting *P. aeruginosa*, *S. haemolyticus*, *A. fumigatus*, and *S. aureus*, enhancing biofilm in the latter (**Figures 1 and 2**). Interestingly, while the early stages of biofilm were affected, attachment itself was unchanged upon addition of either of the three compounds. (**Figure 3**). Both coumarin and umbelliferone altered EPS production, suppressed *pqsA* transcriptional and *pqsR* translational activity, and PYO production in *P. aeruginosa* (**Figures 3**-**5**). Esculetin on the other hand did not influence these phenotypes, suggesting a specific interaction related to the hydroxylation (or lack thereof) at positions 6- and 7- (**Figures 3**-**5**). However, molecular modelling of the coumarin, umbelliferone, and esculetin interaction with the LasR and PqsR receptor proteins did not differentiate their binding affinities or interaction modalities (**Figure 6**). Overall, our computational data suggest that the coumarin derivatives may be able to bind to both PsqR and LasR proteins, but with no significant differences for their binding mode. However, it is worth noting that whereas these coumarin derivatives access the explored binding site at PqsR, access to the binding site of 3-oxo-C12-HSL in LasR is not evident in the ns-time scale of MD simulations.

Some interesting insights also emerge from comparison of the coumarin structures. 4-hydroxy-2-coumarin (**5**) exhibited similarities in biological activity to umbelliferone, suppressing QS in the biosensor assays while enhancing biofilm formation in *S. aureus* NCDO949 and inhibiting biofilm formation in *S. haemolyticus* CUH-T (**Fig. 5**). The position of the hydroxy group (4-hydroxy) and its proximity to the 6-hydroxy group may be relevant here. A methoxy group at the 4-position (4-methoxy-2-coumarin) seems to be well tolerated (compounds **6** and **9**) resulted in strong inhibition of *A. fumigatus* Af293 biofilm formation. Conversely, a hydroxyl group at the same position appears to abrogate activity (compounds **5** and **10**). 7-amino-4-(trifluoromethyl)-2-coumarin (compound **8**) was a strong inhibitor of *Staphylococcus* species biofilm formation, being also active against *A. fumigatus* Af293. Stuctural insights at the molecular level should also be viewed in context of the recently reported diversification of the PqsR protein, being amongst the most variable of the LTTR proteins encoded in *P. aeruginosa* (62). Fine tuning of signal production in bacterial pathogens may offer a more precise approach to infection control.

The evidence from this study would indicate that coumarins and structurally similar compounds can be either ‘bad communicators’ or ‘coercives’ in nature, depending on dose and the molecules-species in question. ‘Bad communicators’ can be considered to have no direct implication on the integrity of the receiver cells, they are not harming the cells but are scrambling the communication i.e., the QSI molecules. ‘Coercive’ molecules negatively impact the receiver cells as is the case of bactericidal antibiotics which prevents the cells achieving minimal signal threshold (17). The data presented here would suggest that the anti-virulence activities of coumarins occurs largely as a consequence of growth-independent alterations to cell behaviour. In some ways, it could be viewed in the same light as the antimicrobial as a weapon or signal (63–65); dose is key to contextualising the outcome. When one considers the distinct coumarin structures exhibiting activity against pathogens that independently carry the LuxIR-AHL, or LuxS, or AIP systems, it could be reasoned that the anti-infective activity of coumarins will be underpinned by interactions outside of these classical systems. It is important that these complexities are deciphered, particularly where coumarins are emerging as significant signals within the complex polymicrobial dynamics of natural communities.

The production of coumarin compounds by a range of plant species may also offer a novel mechanism for control of opportunistic pathogens, whereby the profile of coumarin compounds introduced by a plant-rich diet, may lead to moderation of pathogenic species within communities; the dynamics of the rhizosphere introduced into a host-context (66). While this would mean that coumarin compounds would need to remain stable during passage through the host, and would need to be refractive to biotransformation, it is worth exploring the possibility that coumarin compounds could themselves act as behavioural modulators in human pathogenic species, both fungal and bacterial. Some evidence of a role for the gut microbiome in biotransformation of coumarins has already been reported (67). However, a greater understanding of the ecology of microbial communities and the nature of the small molecule interactions governing their dynamics at a polymicrobial or community level is warranted, such that better and more precise interventions can be designed.

## Materials and Methods

### Bacteria and culture conditions

*Pseudomonas aeruginosa* and *Escherichia coli* were grown in Luria-Bertani (LB) media while *Staphylococcus aureus* and *Staphylococcus haemolyticus* were grown in Tryptic Soy media (TSA or TSB). All bacteria were incubated while shaking at 150 rpm at 37°C. *Aspergillus fumigatus* was routinely grown on *Aspergillus* Minimal Medium (AMM) agar and spores were collected in PBS supplemented with 0.05% (v/v) Tween 20. Coumarin compounds used in this study were dissolved in Dimethyl Sulfoxide (DMSO) and stored at 4°C.

### Quorum-Sensing inhibition assays

*Serratia marcescens* SP19, *Chromobacterium violaceum* DSM30191, and *Agrobacterium tumefaciens* NTL4 were used to investigate quorum sensing inhibition activity against AHLs of varied chain lengths respectively (68–70). *S. marcescens* SP19 and *C. violaceum* DSM30191 were spread over LB agar plates at an OD_600nm_ of 0.5 using cotton swabs. Wells were punctured into the agar plates and the compounds were added at 200 μg, 150 μg, 100 μg or 50 μg. *A. tumefaciens* NTL4 was inoculated into agar at an OD_600nm_ of 0.5 and supplemented with 50 μg ml^−1^ of 5-bromo-4-chloro-3-indolyl-β-d-galactopyranoside. Aliquots of 80 μl were then supplemented with 20 μl of 100 μM 3-oxo-C12-HSL and added to 96-well plates. Coumarins were added at 200 μg, 150 μg, 100 μg or 50 μg. All biosensor strains were incubated at 30°C for 24 hrs and the effect on colour and growth were documented. Phosphate buffered saline (PBS) acted as the control. To assess metabolic activity in the zones of inhibition, XTT (2,3-bis-(2-methoxy-4-nitro-5-sulfophenyl)-2H-tetrazolium-5-carboxanilide) was added in 20 μl aliquots to the zones of inhibition on the QS biosensor agar plates and incubated for 20 mins at room temperature as described previously (71). Metabolic activity was qualitatively assessed through the presence of a red pigment surrounding the test wells.

### Biofilm Assays

#### Microtitre based biofilm assays

Overnight cultures were prepared in fresh media at OD_600nm_ 0.05 and were transferred into 96-well plates (Cellstar, Sigma Aldrich) with the respective coumarin treatments. Plates were grown statically overnight. The liquid cultures were removed following incubation, and the plates were rinsed with sterile deionised water. The plates were allowed to air dry for 20 min. Wells were stained with 200 μl of 0.1% (w/v) crystal violet for 30 min and then washed in deionised water to remove unbound crystal violet. The bound stain was eluted by addition of 200 μl of 96% ethanol. Biofilm formation was monitored and quantified after 20 min by measuring the absorbance at Abs_595nm_ with a Thermo Scientific™ Multiskan™ GO Microplate Spectrophotometer. Time course biofilm assays were performed as above with each time point being prepared as a single plate. Plates were processed as above at 2, 4, 6, 8, and 10 hrs.

#### MBIC Biofilm Assays

MBIC biofilm assays were carried out in standard 96 well plates. *P. aeruginosa* PA14 was inoculated overnight in LB at 37°C shaking. The wells were first filled with 100 μl LB media and compounds were added at a concentration of 32 mM in 100 μl. Serial dilution was carried creating a range from 8 mM to 15.625 μM. The cultures were added in 100 μl volumes at a starting OD_600nm_ of 0.05, to a total volume of 200 μl in each well. Plates were incubated overnight for 20 hrs at 37°C, after which all liquid was removed from the wells, and they were rinsed with water. Crystal violet staining and quantification was performed as described above.

#### Attachment assays

Attachment assays were performed as above with the exception that the starting OD_600nm_ of 0.25 was used and plates were incubated for only 2 hrs prior to processing.

#### EPS production assays

Exopolysaccharide (EPS) production was measured on LB media in 6-well plates supplemented with 20 μg/ml of Coomassie Brilliant Blue and 40 μg/ml of Congo red (72,73). Each coumarin compound was added to a final concentration of 2 mM and an inoculum of the overnight culture was spotted on the centre of each well. Plates were incubated for 120 hrs at 37°C and visualised daily as the staining of cells developed. In addition to testing PAO1 and PA14 strains, an additional mutant strain of PA14 sourced from the non-redundant library was included (*tbpA*, *PA14_13660*), based on its constitutive EPS positive phenotype.

### Growth Curve Analysis

Bacteria were inoculated at 0.05 OD_600nm_ in respective liquid media with coumarin compounds at a final concentration of 2 mM. The bacterial growth was monitored in 100-well honeycomb plates on a BioScreen C. Compound **8** (7-amino-4-trifluromethyl-2-coumarin) is a dye, and interference with the automated OD_600nm_ readings led us to adopt a viable cell count protocol. Glass universals containing 5 ml LB with a starting of 0.05, and a concentration of 2 mM compound 10 were incubated at 37°C with 150 rpm shaking. At 4-, 8- and 12 hr time points, dilutions in PBS were plated on LB with DMSO acting as a control.

### Promoter fusion analysis

Cells from overnight *P. aeruginosa* cultures carrying the appropriate promoter fusion were transferred into fresh media at OD_600nm_ 0.02 and incubated with shaking at 37°C. Cells were sampled at intervals to assess promoter activity using the Miller Assay described previously (49). Miller assays performed with compound **8** were conducted using viable cell counts and are presented as Miller Equivalents where log10 (cfu/ml) replaces OD_600nm_ in the calculation.

### Pyocyanin production

Conical flasks containing 25 ml of LB broth were inoculated with *P. aeruginosa* at an OD_600nm_ of 0.05 and supplemented with 2 mM of each compound. LB broth with DMSO and untreated culture were used as controls. Following incubation of 20 hrs at 37°C with shaking at 180 rpm, 5 ml of culture was removed and centrifuged at 5000 g for 7 min. Pyocyanin was then extracted as described previously (74). The concentration (μg/ml) was calculated using the following formula: concentration (μg/ml) = Abs_520nm_ x 17.072 x dilution factor.

### Pseudomonas quinolone signal (PQS) and 2-heptyl-4-quinolone (HHQ) extraction and TLC analysis

Overnight cultures of *P. aeruginosa* PAO1 were transferred at a starting OD_600nm_ of 0.05 to 25 ml of LB media in conical flasks. The compounds are introduced for a final concentration of 2 mM, DMSO and untreated culture acted as controls. The conical flasks were incubated at 37°C at 180 rpm. After 8 hrs, 10 ml of the culture was removed from which an organic extraction of HHQ and PQS was performed and analysed by Thin Layer Chromatography (TLC) according to the protocols established by Fletcher and colleagues (75).

### Molecular modelling of coumarins at the PqsR and LasR receptors

The Cartesian coordinates of the two transcriptional regulators from *P. aeruginosa* PqsR (UNIPROT id. Q9I4X0) and LasR (UNIPROT id. P25084) were obtained from the crystallographic structures of PqsR in complex with triazolo-pyridine inverse agonist A (PDB id. 6YIZ) (42) and LasR in complex with 3-oxo-C12-HSL (PDB id. 2UV0) (43), respectively. A detailed description of the methodology applied to docking and modelling of the coumarins and LasR or PqsR interaction is described in **Supplementary File S14**. Docking studies were performed with a grid of 60 x 60 x 60 Å dimensions (76), using the genetic algorithm (GA) implemented in AutoDock4 (77). These structures were used for further MD simulation studies. The protein residues were described with the AMBER force field parameters ff19SB (78) and the atoms of the small molecules were described as GAFF/AMBER atom types (79–82). All proteins were embedded in a truncated octahedron of water molecules (TIP3P) (83) and two different setups were built: (a) the docking solution as a protein monomer with a small single molecule at the binding site; (b) the protein with 10 (PqsR, protein monomer) or 20 (LasR, as protein dimer) small molecules in the solution but not in the binding site from the beginning. Long-range interactions were calculated using Particle Mesh Ewald summations (84) using Periodic Boundary Conditions and a cutoff of 10 Å for non-bonded interactions. All MD simulations were run using the suite of programs Amber20 (85) and the MD trajectories were analysed using cpptraj v5.1.0 (86). Pocket and tunnel screening was performed using the Caver web server, predicting the first possible pockets in the apo form of both proteins (44). Caver default parameters were used in all cases.

### Statistical analysis

All experiments performed contained minimum three independent biological replicates. Statistical analysis was carried out on the GraphPad software using one-way ANOVA testing followed by Dunnett’s multiple comparison. Differences were considered significant when the p value was ≤0.05 as indicated by an asterisk.

**Table 1:**
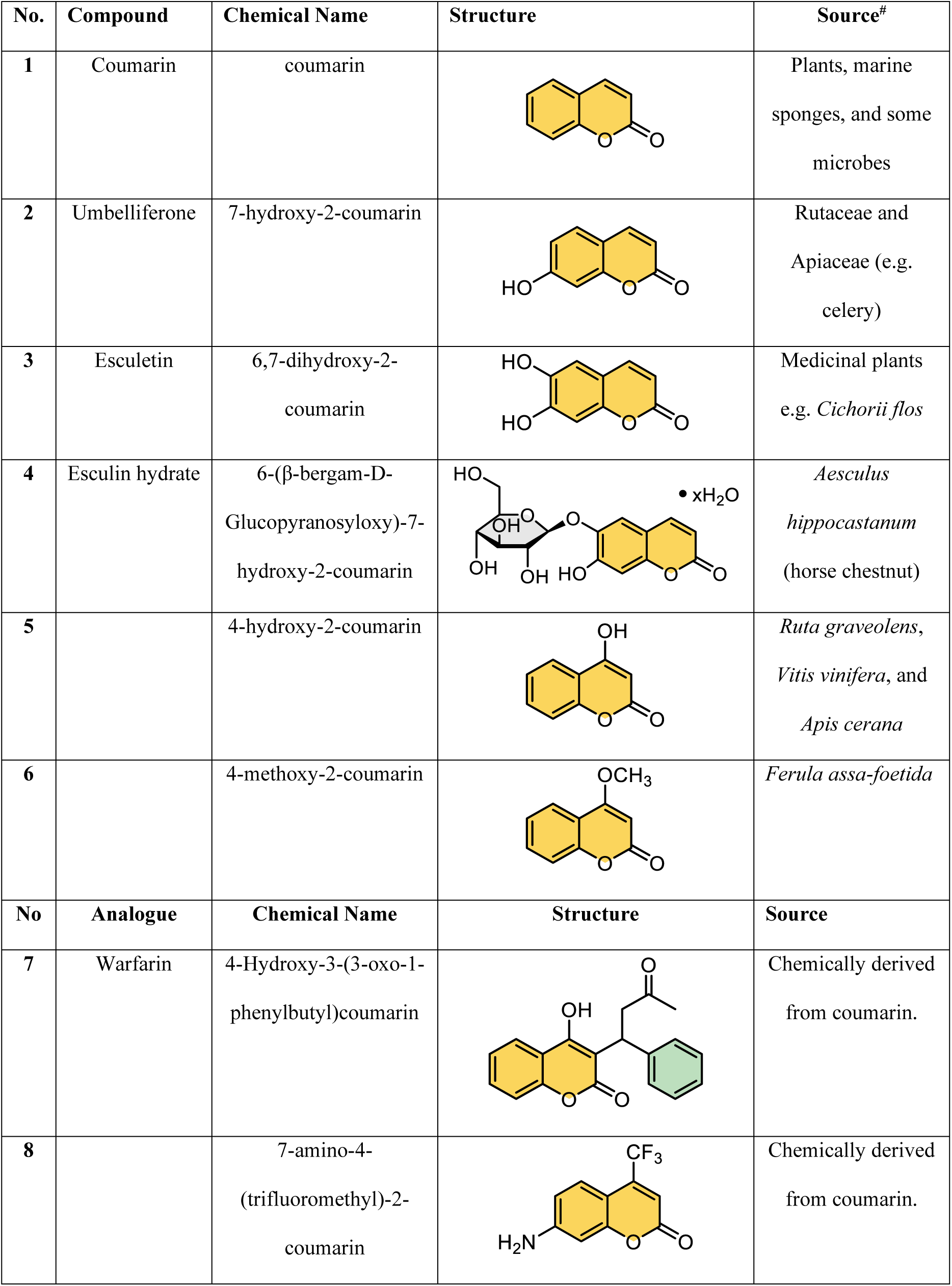
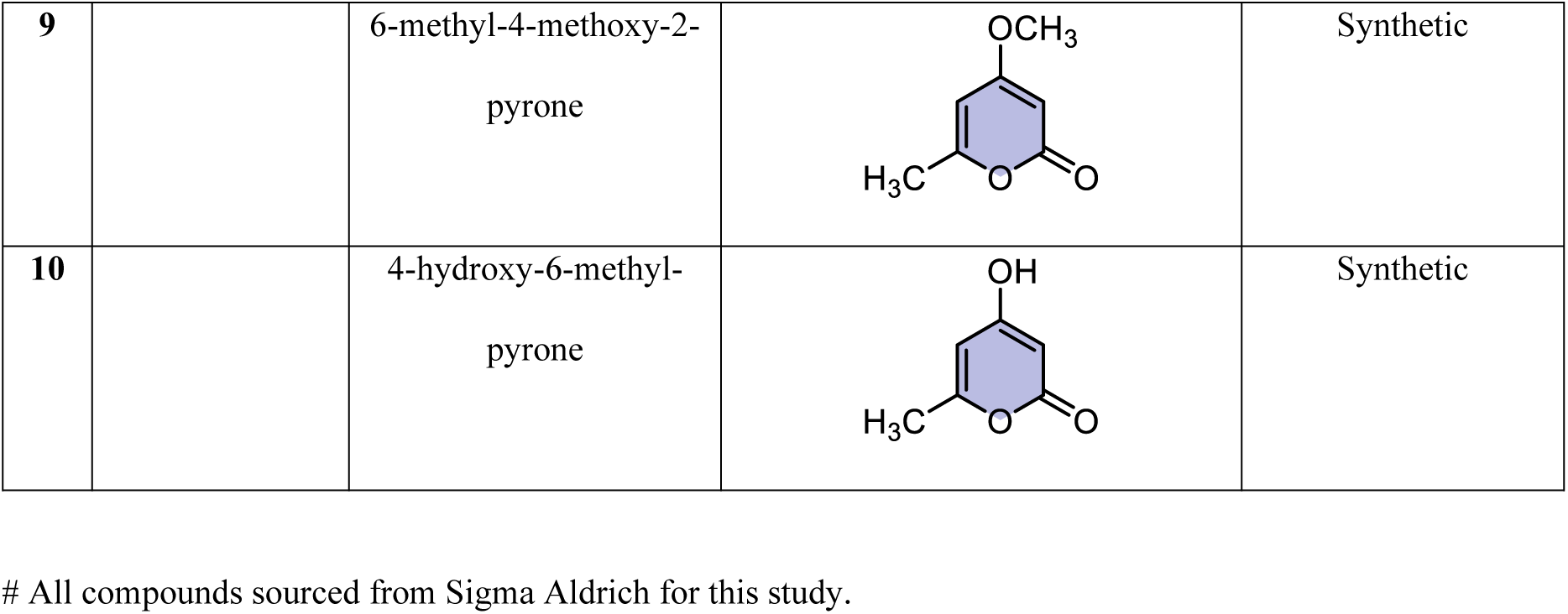
Coumarin structures, sources, and natural sources.

| No. | Compound | Chemical Name | Structure | Source<sup>#</sup> |
| --- | --- | --- | --- | --- |
| 1 | Coumarin | coumarin |  | Plants, marine sponges, and some microbes |
| 2 | Umbelliferone | 7-hydroxy-2-coumarin |  | Rutaceae and Apiaceae (e.g. celery) |
| 3 | Esculetin | 6,7-dihydroxy-2-coumarin |  | Medicinal plants e.g. *Cichorii flos* |
| 4 | Esculin hydrate | 6-(β-bergam-D-Glucopyranosyloxy)-7-hydroxy-2-coumarin |  | *Aesculus hippocastanum* (horse chestnut) |
| 5 |  | 4-hydroxy-2-coumarin |  | *Ruta graveolens*, *Vitis vinifera*, and *Apis cerana* |
| 6 |  | 4-methoxy-2-coumarin |  | *Ferula assa-foetida* |
| No | Analogue | Chemical Name | Structure | Source |
| 7 | Warfarin | 4-Hydroxy-3-(3-oxo-1-phenylbutyl)coumarin |  | Chemically derived from coumarin. |
| 8 |  | 7-amino-4-(trifluoromethyl)-2-coumarin |  | Chemically derived from coumarin. |
| **9** |  | 6-methyl-4-methoxy-2-pyrone |  | Synthetic |
| **10** |  | 4-hydroxy-6-methylpyrone |  | Synthetic |

## Acknowledgements

The authors acknowledge Prof. Ruth Massey and Prof. Ronan McCarthy for careful reading of the manuscript and for the provision of critical insights. The authors also thank Prof. Paul Williams for insightful experimental suggestions. FJR and GMG acknowledge the support of Research Ireland and Science Foundation Ireland (SSPC-3,12/RC/2275_2). FJR acknowledges the support of the Health Research Board (HRB-ILP-POR-2019-004). GMG would also like to acknowledge Research Ireland and Science Foundation Ireland (SFI/12/IP/1315 and 21/FFP-A/8784) and a research infrastructure award for process flow spectroscopy (ProSpect) (grant: SFI 15/RI/3221 and 21/RI/9705). DPR and PASM acknowledge Medical University of Graz for computation time in MedBioNode. MC acknowledges the Irish Research Council Postgraduate Scholarship (GOIPG/2021/692). For the purpose of Open Access, the author has applied a CC BY public copyright licence to any Author Accepted Manuscript version arising from this submission. The funders had no role in study design, data collection and interpretation, or the decision to submit the work for publication

## Data Availability Statement

All data pertaining to the manuscript is contained within the supplementary files, and any additional materials described in the manuscript, including all relevant raw data, will be made freely available to any researcher wishing to use them for non-commercial purposes.

## Author Contributions

Conceptualisation: FJR. Experimentation: DB, BOR, MC, AG, DW, JL, DJPR, PASM, and FJR. Funding: FJR, GmcG, MC, and PASM. Initial Draft of Manuscript: DB, BOR, MC, and FJR. Review and Final Drafts: DB, BOR, MC, AG, DW, JL, DJPR, PASM, GMG, and FJR.

---

**Supplementary Table 1:**
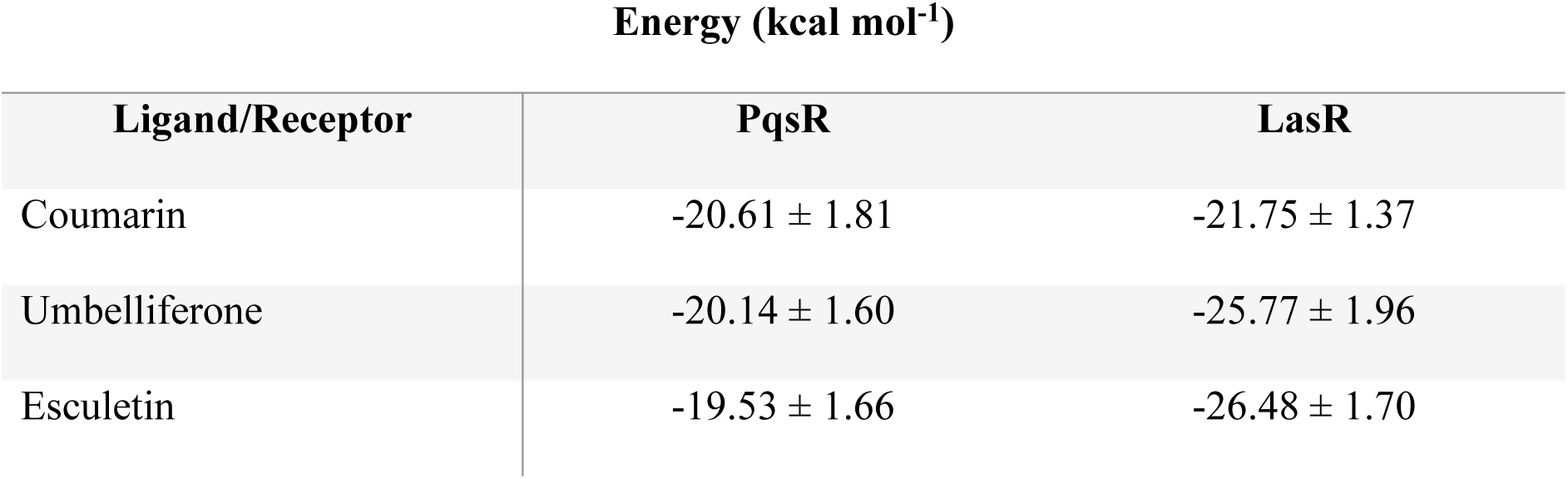
Total ligand-protein interaction energy (kcal mol −1) obtained using MM-ISMSA3 (values are reported as total energy ± standard deviation). A 30ns-window with a total of 30 snapshots were used in each case for the analysis.

**Supplementary Table S2:** Bottleneck radius (°A) and average throughput for the tunnels found by Caver web server.

**PqsR**
| Tunnel | Bottleneck Radius ($^{\circ}$A) | Avg. Throughput |
| --- | --- | --- |
| 1 (blue) | 1.74 | 0.82 |
| 2 (green) | 1.04 | 0.68 |
| 3 (red) | 1.04 | 0.58 |
| LasR |  |  |
| 1 (blue) | 1.41 | 0.74 |
| 2 (green) | 0.94 | 0.56 |
| 3 (red) | 1.13 | 0.49 |

## Supplementary Files

**Supplementary Figure S1.** Visualisation of quorum sensing inhibition (**A**) in the absence and (**B**) presence of XTT for detection of metabolic activity. Coumarin compounds are added at 50 (bottom left), 100 (top left), and 150 (top right) to each well. All data presented is representative of three independent biological replicates. PBS is presented as a control.

**Supplemental Figure S2. Dose dependent biofilm inhibition in ESKAPEE pathogens in the presence of natural coumarin compounds.** (**A**) Summary inhibition of biofilm formation by coumarin compounds at 0.01, 0.1, 1, and 2 mM. (**B**) Inhibition of biofilm formation by 1 mM coumarin compounds. Each dataset is the mean (+/- SEM) of at least three independent biological replicates. Statistical analysis was performed by one-way ANOVA with Dunnett’s post-hoc corrective testing (* p≤0.05).

**Supplementary Figure S3. MBIC analysis of *P. aeruginosa* biofilm formation in the presence of coumarin, umbelliferone, and esculetin.** Data presented is the mean (+/- SEM) of three independent biological replicates.

**Supplementary Figure S4. EPS production in *P. aeruginosa* PA14 in the presence of coumarin compounds**. Colony formation was monitored over 120 hrs and visualised for evidence of EPS production. DMSO was included as a carrier control, with the constitutive EPS producing PA14 TnM *tbpA-D* mutant included for comparison.

**Supplementary Figure S5. Growth kinetics of ESKAPEE pathogens in the presence of 2 mM coumarin compounds.** Test organisms were assayed on a Bioscreen C plate reader. All data presented is the mean (+/- SEM) of three independent biological replicates.

**Supplementary Figure S6. (A)** RMSD (Å) of the 10 small molecules with respect to docking results in the PqsR receptor. Highlighted we show the molecules with high resemblance with the docking results. **(B)** RMSD (Å) of the 20 small molecules with respect to docking results in the LasR receptor. Each row of images corresponds to one chain of the dimer.

**Supplementary Figure S7. Growth kinetics of ESKAPEE pathogens in the presence of 2 mM coumarin analogues.** Test organisms were assayed on a Bioscreen C plate reader. All data presented is the mean (+/- SEM) of three independent biological replicates.

**Supplementary Figure S8.** The effect on growth of (A) CUH-T and (B) NCDO949 in the presence of compound 10 at 2 mM. Due to the incompatibility of using compound 10 with the Bioscreen, the growth was measured by viable cell count plating over a 12 hr period and is expressed as cfu/ml. All data presented is the mean (+/- SEM) of three independent biological replicates.

**Supplementary Figure S9. Biosensor analysis of coumarin analogue structures indicates lack of QS suppression upon modification of the core coumarin.** (i) zone inhibition of pigment production (ii-iii) visualisation of pigment production in biosensor strains.

**Supplemental Figure S10. Biofilm formation in ESKAPEE pathogens and *A. fumigatus* in the presence of coumarin analogues.** Biofilm formation is presented as OD_595nm_, and visualisation of *A. fumigatus* biofilms is presented below the quantitative graphs. Each dataset is the mean (+/- SEM) or is representative (images) of at least three independent biological replicates. Statistical analysis was performed by one-way ANOVA with Dunnett’s post-hoc corrective testing (* p≤0.05, ** p≤0.005, *** p≤0.001).

**Supplemental Figure S11. Dose dependent biofilm formation in ESKAPEE pathogens in the presence of coumarin analogues.** Each dataset is the mean (+/- SEM) of at least three independent biological replicates. Statistical analysis was performed by one-way ANOVA with Dunnett’s post-hoc corrective testing (* p≤0.05, ** p≤0.005, *** p≤0.001).

**Supplementary Figure S12. EPS production in *P. aeruginosa* PA14 in the presence of synthetic coumarin compounds.** All data presented is representative of three independent biological replicates.

**Supplementary Figure S13. Quorum sensing promoter fusion analysis in *P. aeruginosa*. (A-C)** The three systems operating in *P. aeruginosa* were studied in the presence of synthetic coumarin compounds. All data presented is the mean (+/- SEM) of at least three independent biological replicates. **(D-G)** Analysis of promoter activity in response to compound **8** was performed using viable cell counts rather than OD_600nm_.

